# Identification of SARS-CoV-2 Antiviral Compounds by Screening for Small Molecule Inhibitors of the nsp14 RNA Cap Methyltransferase

**DOI:** 10.1101/2021.04.07.438810

**Authors:** Souradeep Basu, Tiffany Mak, Rachel Ulferts, Mary Wu, Tom Deegan, Ryo Fujisawa, Kang Wei Tan, Chew Theng Lim, Clovis Basier, Berta Canal, Joseph F. Curran, Lucy Drury, Allison W. McClure, Emma L. Roberts, Florian Weissmann, Theresa U. Zeisner, Rupert Beale, Victoria H. Cowling, Michael Howell, Karim Labib, John F.X. Diffley

**Author notes:** These authors contributed equally.

## Abstract

The COVID-19 pandemic has presented itself as one of the most critical public health challenges of the century, with SARS-CoV-2 being the third member of the *Coronaviridae* family to cause fatal disease in humans. There is currently only one antiviral compound, remdesivir, that can be used for the treatment of COVID-19. In order to identify additional potential therapeutics, we investigated the enzymatic proteins encoded in the SARS-CoV-2 genome. In this study, we focussed on the viral RNA cap methyltransferases, which play a key role in enabling viral protein translation and facilitating viral escape from the immune system. We expressed and purified both the guanine-N7 methyltransferase nsp14, and the nsp16 2’-O-methyltransferase with its activating cofactor, nsp10. We performed an *in vitro* high-throughput screen for inhibitors of nsp14 using a custom compound library of over 5,000 pharmaceutical compounds that have previously been characterised in either clinical or basic research. We identified 4 compounds as potential inhibitors of nsp14, all of which also show antiviral capacity in a cell based model of SARS-CoV-2 infection. Three of the 4 compounds also exhibited synergistic effects on viral replication with remdesivir.

## Introduction

The SARS-CoV-2 virus is a novel respiratory pathogen that is able to infect both animals and humans, resulting in the disease COVID-19, which was officially declared a global pandemic by the World Health Organisation (WHO) on 11^th^ March, 2020 [1]. This is the third novel zoonotic virus belonging to the genus *Betacoronavirus* since the start of the 21^st^ century, after the original SARS-CoV-1 in 2003, and MERS-CoV in 2012 [2]. What sets SARS-CoV-2 apart from the previous two is its distinctive virulence on an overall population, as well as its epidemiological dynamics (reviewed in [3]). Therefore, vastly different detection and containment strategy have been required compared to those that were used successfully with SARS-CoV-1 and MERS-CoV outbreaks.

SARS-CoV-2 is a membrane enveloped virus with peplomer-forming spike (S) glycoproteins on its surface, giving it the characteristic “corona” shape when visualised under electron-microscopy (EM) [4]. The structural components of the virus are highly variable and mutable, with over 10% of the open reading frame mutations identified between December 2019 to April 2020 found to occur in the Spike glycoprotein gene alone [5]. The highly mutative nature of the viral coat therefore poses a problem for producing effective long-term neutralising antibodies through administration of vaccines, leading to viral strains such as B1.351 that escape from the immune response [6].

However, the structural proteins of the virus actually only constitute a small part of the coding capacity of the viral genome, with only 4 out of 29 proteins encoding viral structural components (Figure 1A) [7]. A pair of very large open reading frames (Orf1a and Orf1ab) comprise the first two-thirds of the genome. The encoded polyproteins (pp1a and pp1ab) are autoproteolytically cleaved into 16 distinct non-structural proteins (nsp), generating the enzymes and accessory proteins responsible for viral replication once inside a eukaryotic cell.

**Figure 1:**
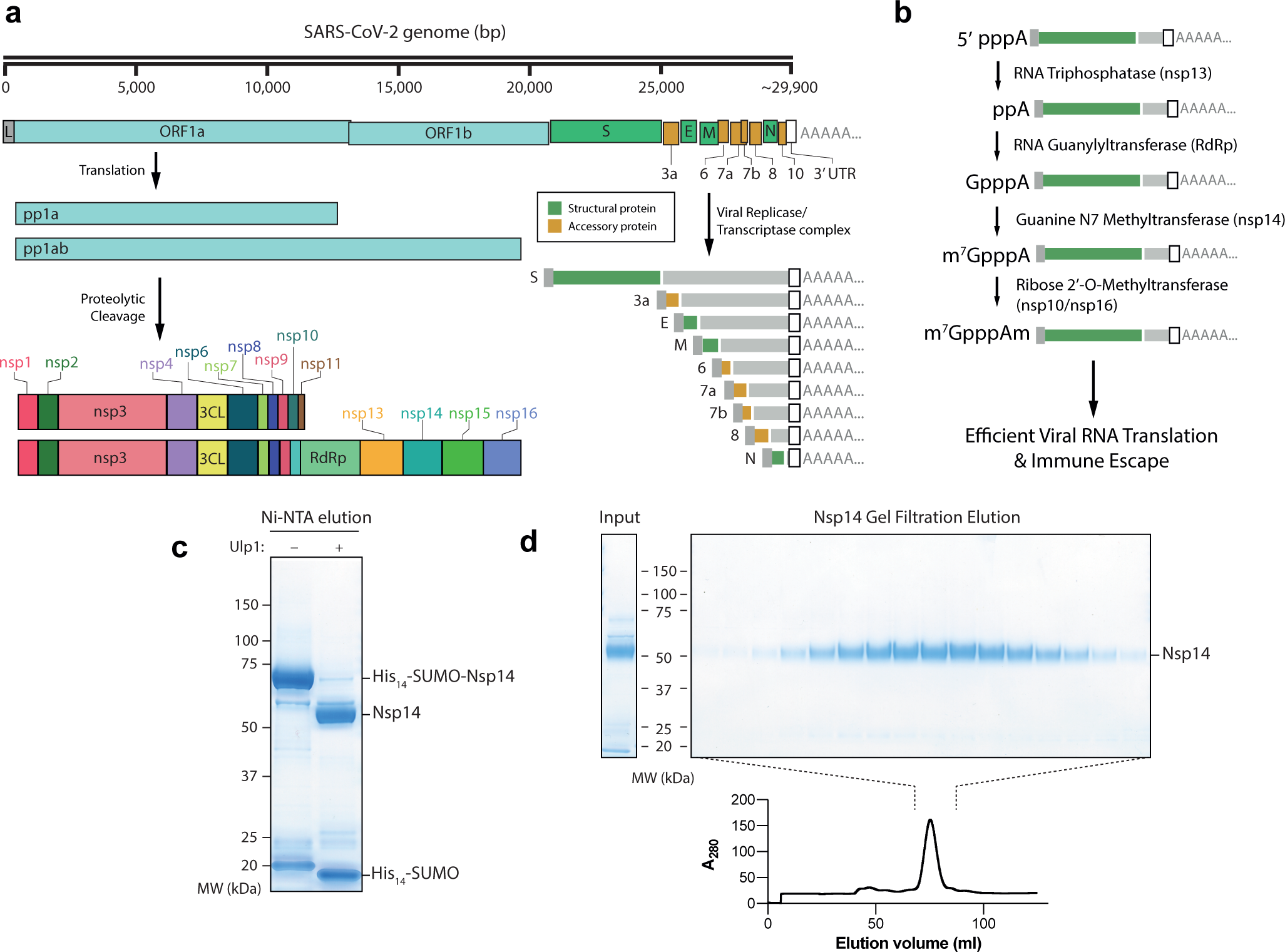
Purification of nsp14 Guanine N7 Methyltransferase. **a)** Outline of the SARS-CoV-2 genome. Pp1a and Pp1ab represent polyproteins a and ab, respectively. Pp1a and pp1ab are able to autoproteolytically cleave themselves to form the nsp proteins outlined. The viral replicase/transcriptase complex produces a series of nested viral RNAs that encode accessory (orange) or structural (green) viral proteins. **b)** Viral RNA capping outline. The initial RNA nucleotide possesses a γ and β phosphate, unlike following RNA bases. The γ phosphate is removed by nsp13, followed by the addition of Gp by nsp12, releasing pyrophosphate. nsp14 and nsp16/10 then catalyse the formation of the final Cap-0 structure. **c)** Coomassie gel of His_14_-SUMO cleavage. Left column: Elution from Ni-NTA beads without the Ulp1 SUMO-dependent protease. Right: Elution from Ni-NTA beads after treatment with Ulp1 (see Methods). **d)** Gel filtration fractions of nsp14. Left: Input to gel filtration. Right: Pooled fractions from the main peak of the elution (lower). nsp14 expected size: 55 kDa.

Coronavirus RNAs are produced by the viral replicase/transcriptase complex, and are post transcriptionally capped at the 5’ end with a cap structure similar to endogenous mRNA caps, which are necessary for efficient translation and RNA stability [8–11]. Crucially, viral RNA capping has been shown to be essential for the synthesis of viral proteins through eukaryotic translation initiation factor 4E (eIF4E) recognition [12, 13]. In addition, RNA cap methylation is also important for ensuring efficient ribosome binding and engagement of the host translation machinery, as well as avoiding degradation by exoribonucleases [9, 14].

Aside from reduced translational capacity, uncapped RNA also triggers the host innate immune response leading to the expression of antiviral cytokines, which limit virus replication and shape adaptive immunity [15]. Other host sensor proteins recognise incomplete or absent cap RNA structures on viral RNA, and are responsible for the inhibition of viral translation [16, 17]. Formation of the viral RNA cap structures thus protect the virus against cell intrinsic antiviral effectors and potentiate the infective potential of the virus. For these reasons, viral enzymes involved in the capping the viral RNA are promising targets for antiviral drug development [18–20].

In eukaryotes, these methylated cap structures are added co-transcriptionally upon transcription by RNA Pol II in the nucleus (reviewed in [21]). Because coronavirus replication and transcription occur in the cytoplasm, independently of Pol II, the formation of these cap structures must be catalysed by viral enzymes. Over one-third (6/15) of the coronavirus nsps are necessary to efficiently cap viral RNAs, which exemplifies the complexity of this process as well as its importance for the viral lifecycle. Firstly, following RNA synthesis, nsp13 removes the terminal γ-phosphate from the initiating adenosine nucleotide triphosphate (Figure 1B). Nsp12 then acts as an RNA-guanylyltransferase, generating GpppA-capped RNA (Figure 1B). Subsequently, nsp14 transfers a methyl group to the N7 position of the terminal cap guanine forming the m7G cap structure, otherwise known as cap-0. Finally, the nsp16/nsp10 complex is responsible for 2’-O methylation of the cap ribose to form cap-1, which is the terminal cap structure on viral RNAs (Figure 1B).

The nsp14 enzyme in SARS-CoV-2, similar to other coronaviruses, carries dual functionality as both a methyltransferase and a 3’-5’ exoribonuclease. While the exoribonuclease activity is dependent upon the nsp10 cofactor, this cofactor is not required for the N7-MTase function in nsp14 [18, 22]. In contrast, the other methyltransferase enzyme, nsp16, which converts cap-0 to cap-1, requires the presence of nsp10 to act as an allosteric activator to increase RNA substrate binding affinity [22, 23]. Both nsp14 and nsp16/10 are dependent upon the presence of S-adenosyl-L-methionine (SAM), which acts as the methyl donor for the respective methylated cap modifications, and S-adenosyl-L-homocysteine (SAH) is the resulting by-product. Using this property, we adopted a biochemical assay that detects the conversion of SAM to SAH to measure the relative methyltransferase activity. We then proceeded to first investigate the *in vitro* methyltransferase activity of purified nsp14 enzyme, and screened against a custom library of over 5000 characterised chemical compounds, with the aim of identifying potential antiviral drugs.

## Results

### Expression & purification of nsp14

To generate untagged nsp14 we used the His-SUMO tag [24], which can be completely removed by the SUMO protease, Ulp1. His_14_-SUMO-nsp14 was expressed in *Escherichia coli* cells after induction with IPTG overnight. Clarified cell extract was then passed over a Ni-NTA column and His_14_-SUMO-nsp14 was eluted with imidazole (Figure 1C). The His_14_-SUMO tag was cleaved by addition of the Ulp1 (Figure 1C) and was further purified by gel filtration chromatography, where untagged nsp14 eluted as a single peak (Figure 1D). This methodology allowed the expression and subsequent purification of nsp14 in high yield.

To ensure that purified nsp14 was functional, we examined the ability of the enzyme to catalyse methylation of the G(5’)pppG(5’) cap in a capped RNA substrate. Radiolabelled ^32^P-RNA substrate was incubated with nsp14, the viral nsp16 methyltransferase, human CMTR1 (Cap-specific mRNA 2’-O-Methyltransferase 1) and RNMT (RNA Guanine-N7 Methyltransferase) in complex with its activating co-factor RAM (RNMT Activating Miniprotein). Whilst nsp14 and RNMT-RAM are guanine-N7 methyltransferases that use GpppG-RNA as a substrate [25], CMTR1 and viral nsp16 are 2’-O-methyltransferases that utilise previously guanine-N7 methylated me_7_GpppN-RNA as a substrate. Following incubation with methyltransferase, RNA substrate was digested with Nuclease-P1, which digests RNA but leaves RNA cap structures intact. This digested mixture was then analysed by thin layer chromatography. As expected nsp16 and CMTR1 failed to utilise the GpppG-RNA substrate, whereas both viral nsp14 and human RNMT-RAM efficiently methylated the GpppG-RNA to form me_7_GpppG-RNA (Figure 2A). To confirm that the methylation was strictly dependent on the catalytic activity of nsp14 we mutated aspartate 331, which resides in the catalytic core of nsp14 and is predicted to be essential for methyltransferase activity, to alanine (D331A). The purified nsp14^D331A^ protein was inactive as a methyltransferase in this assay (Figure 2B), confirming that the nsp14 of SARS-CoV-2 functions as a methyltransferase in accordance with the function of nsp14 enzymes from other coronaviruses [26], and demonstrating that our purified wild-type nsp14 is catalytically active.

**Figure 2:**
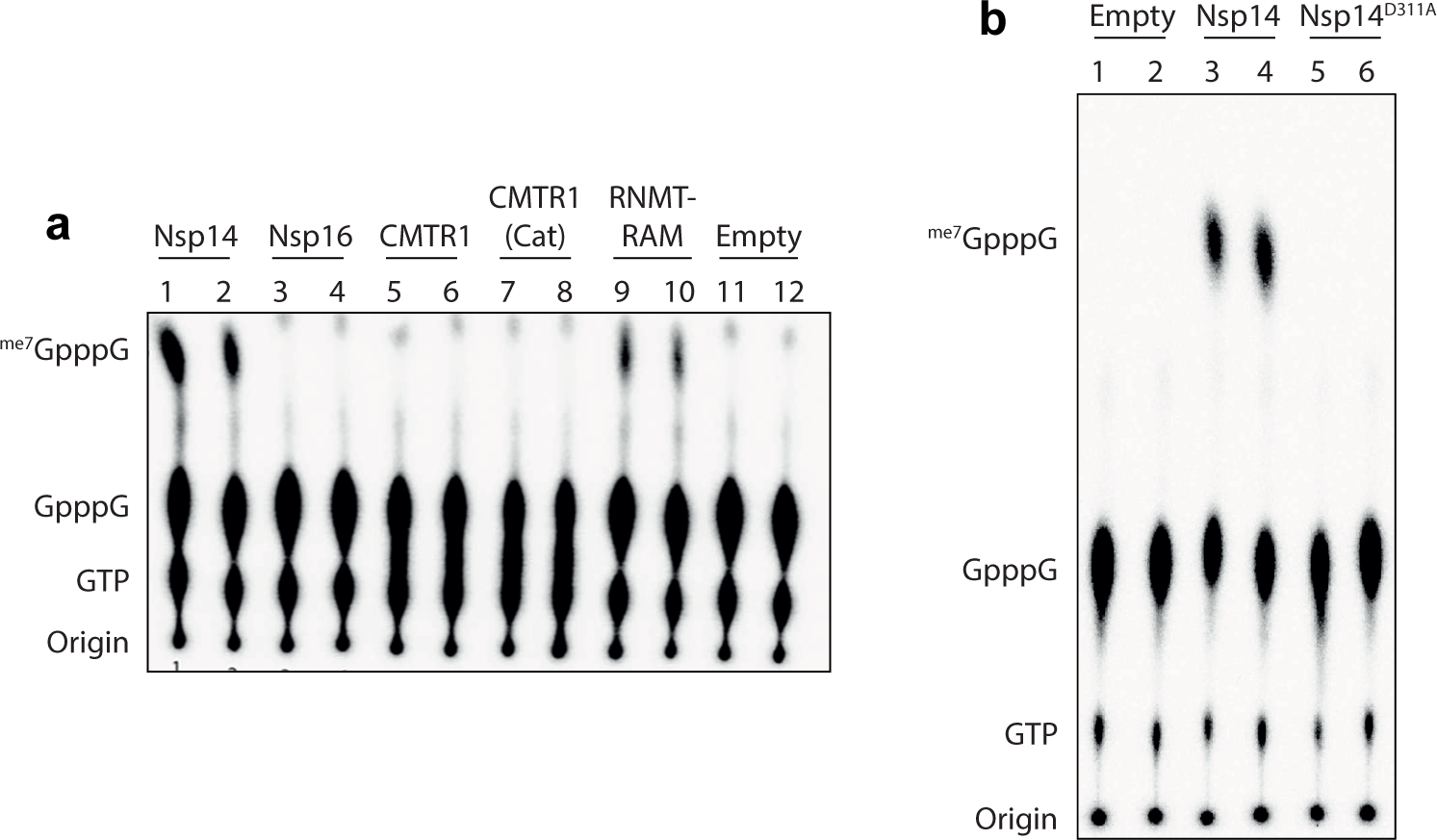
Nsp14 is functional as a cap methyltransferase *in vitro*. **a)** Nsp14, nsp16, CMRT1, the catalytic domain of CMRT1 (CMRT1-Cat), and RNTM fused to the first 80 amino acids of RAM were incubated with radiolabelled GpppG-RNA and the methyl donor S-adenosyl methionine. After the reaction was stopped, RNAs were digested with Nuclease-P1 and resolved by thin layer chromatography (see methods). A negative control was also run in which no enzyme was added. Standards were visualized by UV light to establish correct migration. **b)** nsp14 and nsp14^D331A^ were incubated with radiolabelled GpppG as above, with a negative control given in which no enzyme was added.

### An HTRF based assay for methyltransferase activity

Thin layer chromatography is not amenable to high-throughput screening for nsp14 inhibitors, and therefore we turned to a fluorescence-based readout of methyltransferase activity. We employed a commercially available homologous time-resolved fluorescence (HTRF) assay, which monitors the activity of S-adenosyl methionine (SAM)-dependent methyltransferases. SAM acts as a methyl donor for methyltransferases, yielding S-adenosyl homocysteine (SAH) as a final product (Figure 3A, left). The HTRF assay revolves around a Terbium (Tb) cryptate conjugated to the Fc region of an anti-SAH antibody. This antibody is able to bind SAH which has been conjugated to the d2 fluorophore (SAH-d2) through its variable (Fab) region. In a system without newly produced SAH, the Tb cryptate is able to form a FRET pair with the d2 fluorophore present on the bound SAH-d2 (Figure 3A, right upper). The production of SAH from successful methyltransferase reactions competes with and displaces SAH-d2 from the variable region of the antibody, causing a reduction in signal (Figure 3A, right lower).

**Figure 3:**
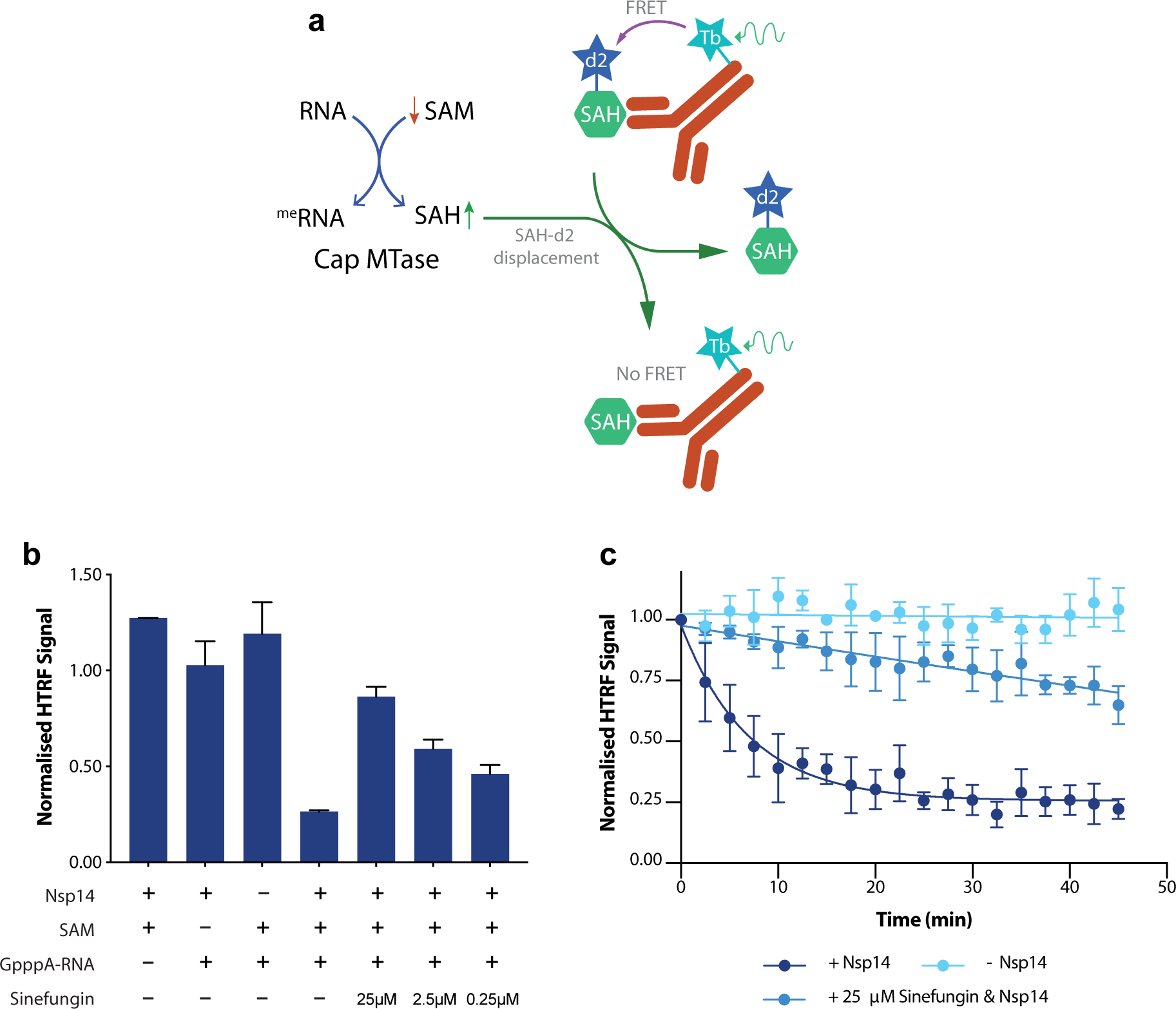
An HTRF based assay for methyltransferase activity. **a)** Outline of the HTRF based assay for methyltransferase activity. Both nsp14 and nsp10-16 are SAM dependent methyltransferases that produce SAH following successful methyltransfer to their substrate. This SAH displaces SAH-d2 from the variable region of an α-SAH Tb cryptate-conjugated antibody, thus lowering HTRF signal through the disruption of the Tb cryptate – d2 FRET pair. **b)** nsp14 was assayed for methyltransferase activity through the HTRF based assay. The methyltransferase reaction was run in either the absence of 10nM nsp14, 1 μM SAM, 0.11 mM GpppA-RNA, or in the presence of all three components. In addition, the methyltransferase reaction was conducted in the presence of 0.25 μM, 2.5 μM and 25 μM of the pan-methyltransferase inhibitor Sinefungin, which acts as a competitive inhibitor (with respect to SAM) towards SAM-dependent methyltransferases. **c)** Time course of nsp14 activity by HTRF assay. 20 nM of nsp14 was incubated with 0.11 mM GpppA cap analogue and 1 μM SAM for the time indicated. In addition, an experiment was run in the absence of nsp14, and in the presence of the methyltransferase inhibitor sinefungin. The reaction was started as a master mix with the addition of nsp14, and 8 μl was removed at every time point and added to 2 μl of 5M NaCl to stop the methyltransferase reaction. Points are the mean of three technical repeats, and error bars indicate range.

A robust reduction in HTRF signal was seen after incubation of nsp14, SAM and GpppA-RNA, which was lost if any one of the three components necessary for methyltransfer was omitted (Figure 3B). SAM dependent methyltransferases are susceptible to inhibition by sinefungin, which acts as a competitive inhibitor (with respect to SAM) of methyltransferases. Figure 3B shows that sinefungin inhibited nsp14 activity in a dose-dependent manner. Monitoring over time, the reaction reaches a plateau after around 20 min (Figure 3C). This plateau is unlikely to be due to time-dependent enzyme inactivation since we observed a shallow but continuous decrease of HTRF signal over the course of a 45-minute reaction in the presence of sinefungin (Figure 3C). Therefore, the rapid plateau reached likely represents a lower boundary for the HTRF assay under our experimental conditions.

We next characterised the range of substrates that nsp14 is capable of methylating. Unlike other guanine-N7 methyltransferases, coronavirus nsp14 has previously been reported to be able to methylate free GpppA cap analogue without attached RNA, as well as free nucleotide GTP [25]. When titrating GpppA-capped RNA, GpppA cap analogue, or nucleotide GTP, we were able to see a reduction in HTRF signal, indicating that SARS-CoV-2 nsp14 retains this unusually broad substrate repertoire, and is able to catalyse methyltransfer to all three substrates (Figure 4A). nsp14 is, however, unable to catalyse methyltransfer to already guanine-N7 methylated me_7_GpppA cap analogue (Figure 4A). This suggests that, despite wide substrate specificity, SARS-CoV-2 nsp14 acts specifically as a guanine-N7 directed methyltransferase. Following this, we were able to quantify the Michaelis constants (K_m_) for all of the known substrates of the nsp14 methyltransferase. nsp14 showed similar K_m_ values for both GpppA and GpppA-RNA substrates (Figure 4B,C), but demonstrated slightly reduced affinity towards nucleotide GTP (Figure 4D).

**Figure 4:**
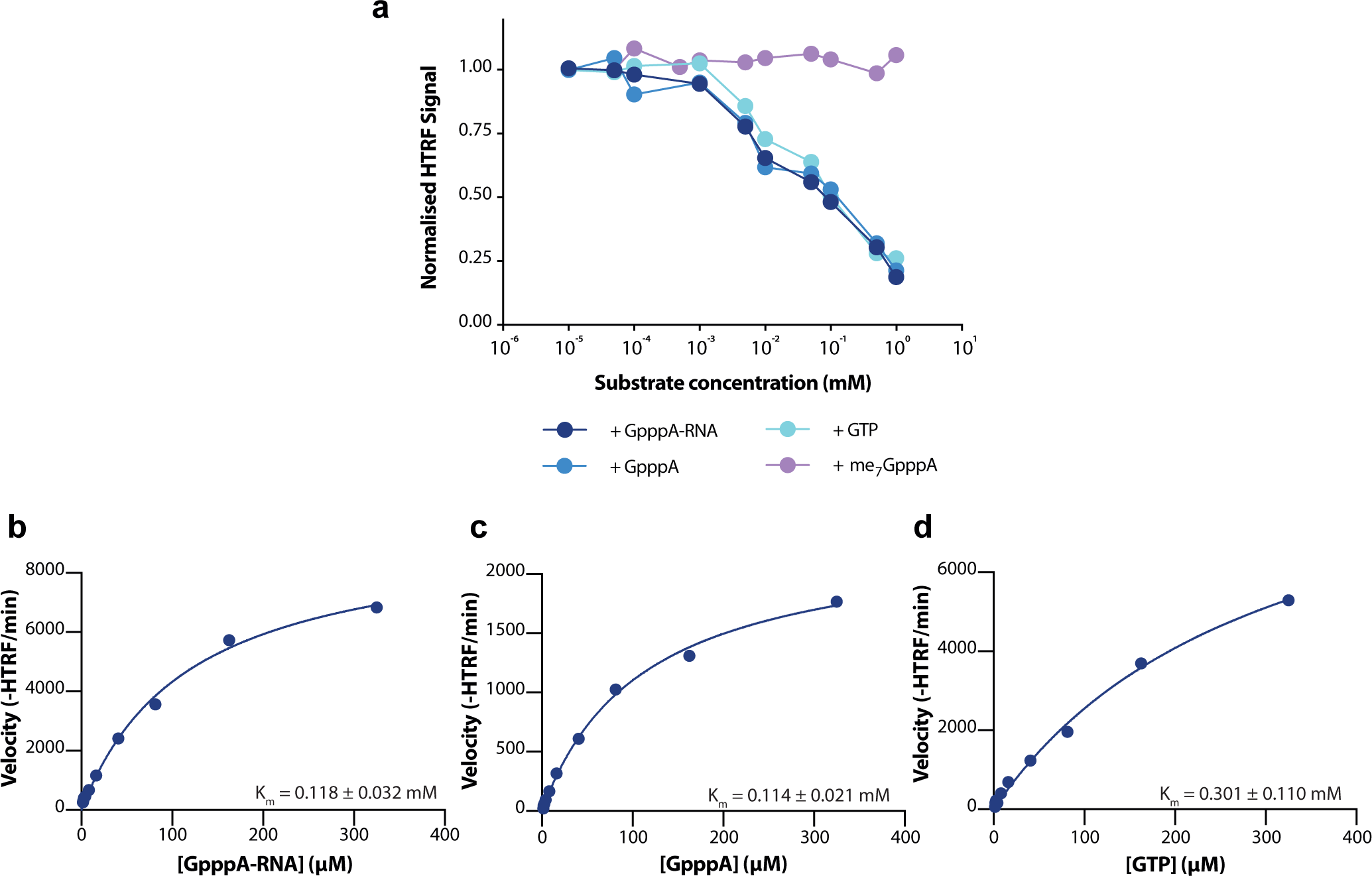
K_m_ values for substrates of nsp14. **a)** Titration of various nsp14 susbstrates. GTP, GpppA cap analogue, GpppA-RNA, and me_7_GpppA cap analogue were incubated for 20 minutes with 5 nM nsp14 in the presence of 1 μM SAM. **b-d)** Determination of Michaelis constants for (**b**) GpppA-RNA, (**c**) GpppA, and (**d**) GTP for nsp14. nsp14 concentration was fixed at 5 nM for all experiments, and substrate concentration varied. V_max_ values cannot be compared between substrates, and are therefore not given. HTRF values are given as raw, and not normalised, HTRF values. Errors given are 95% confidence ranges.

### Primary identification of nsp14 inhibitors

Next, we adapted the HTRF assay conditions to conduct a high-throughput screen against a custom compound library to discover novel nsp14 inhibitors. Primary screening for inhibitors of nsp14 was conducted using a custom compound library containing over 5000 compounds at a concentration of 3.125 μM. Given that nsp14 showed similar K_m_ values between the GpppA-RNA substrate and GpppA cap analogue, we conducted the screen using the cap analogue substrate which did not add the complications of working with an RNA substrate. Library compounds were resuspended in DMSO, and negative controls wells also included DMSO to a final concentration of 0.03125 % (v/v). Sinefungin at 3.125 μM was included in several wells on each plate to serve as a positive control, thereby allowing determination of screen quality.

After screening, we calculated the Z’ factor of our screen to be 0.625, indicating a high-quality screen (Figure S1). We therefore calculated Z-scores for all compounds, and ranked them by increasing Z-score (Figure 5A). We firstly selected ‘hit’ compounds from the list of compounds with a Z-score of above 3.0, but made exceptions for some compounds if their Z-score was >2.5 and either clinically relevant, or a previously characterised methyltransferase inhibitor. This left 83 compounds, which was narrowed by removing compounds which likely represented screening errors (see methods), resulting in a list of 63 hits. These hits, and their associated Z-scores, are listed in Table S1. We subsequently selected compounds to take forward for validation based on if they were available to purchase commercially, if they were rapidly available, if they were known to be tolerated in humans, and favoured particularly those that were already licenced therapeutics. This was done to focus our hits to compounds that may be of clinical relevance to the treatment of COVID-19, and resulted in a list of 15 compounds to carry forward for validation (Table S1).

**Figure 5:**
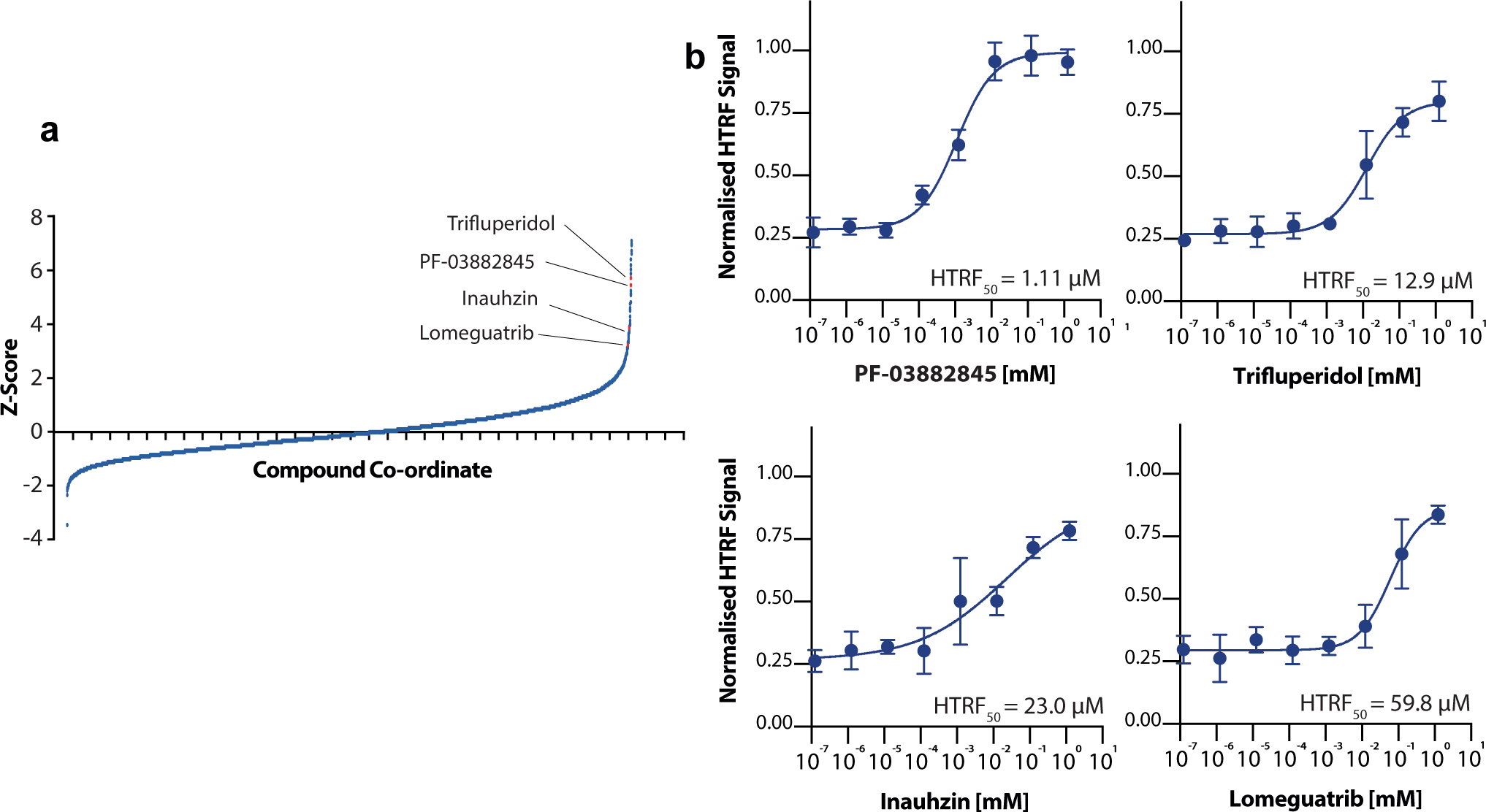
Screening for inhibitors of nsp14. **a)** Z-values of all compounds in the high throughput screen. Compounds are ranked by observed Z-value. Four compounds highlighted were validated in further HTRF experiments. **b)** Titration of validated compounds from high throughput screening. Normalised HTRF_50_ values for each compound are given within the respective panel. Normalised HTRF_50_ values are determined from fitting four-parameter agonist vs. response curves (see Methods).

Using the HTRF-based assay, we found that 4 of these 15 compounds (PF-03882845, Inauhzin, Lomeguatrib, and Trifluperidol) consistently inhibited nsp14 when assayed individually over a wider range of concentrations (Figure 5B and S2). Among these compounds, PF-03882845 was the most potent inhibitor (HTRF_50_ = 1.1 μM), followed by Trifluperidol (HTRF_50_ = 12.9 μM), Inauhzin (HTRF_50_ = 23.0 μM), and finally Lomeguatrib (HTRF_50_ = 59.8 μM). The remaining 11 compounds did not show significant inhibition, and were excluded from further analysis (Figure S2, Table S1).

To test the specificity of the four validated compounds towards nsp14 methyltransferase activity, we checked for cross-inhibition of the other viral methyltransferase nsp16/nsp10. To test the role of these 4 compounds on the nsp16 methyltransferase activity, we purified nsp16 fused to its cofactor nsp10. Fusion of nsp14 to its cofactor nsp10 has been shown an effective strategy to obtain active recombinant nsp14 exonuclease (Canal et al, this issue). Following this strategy, we decided to fuse the methyltransferase nsp16 with its cofactor nsp10 that, similarly to for nsp14, would ensure stoichiometric expression of both subunits as well as their association. In the nsp10-16 fusion protein, nsp10 was placed in the N-terminus followed by the linker GSGSGS and nsp16, and the protein was purified using an N-terminus His_14_-SUMO tag. We purified the nsp10-16 fusion protein to homogeneity from *E. coli* cells in a manner similar to nsp14 (Figure 6A, see methods). We then confirmed that the nsp10-16 fusion 2’-O-methyltransferase was able to give a dose-dependent reduction in HTRF signal when catalysing methyltransfer to its endogenous substrate me_7_GpppA-RNA (Figure 6B). Nsp10-16 was able to methylate me_7_GpppA-RNA efficiently at substantially lower substrate concentrations than nsp14, and demonstrated a Km value ∼100x lower for me_7_GpppA-RNA than nsp14 for GpppA-RNA (Figure 6C). Finally, we then used the nsp10-16 methyltransferase assay to test specificity of our 4 validated compounds towards the inhibition of nsp14 methyltransferase activity. When using this assay, none of the nsp14 inhibitors were able to inhibit nsp10-16, however, the SAM-competitor sinefungin was able to inhibit methyltransfer by nsp10-16 (Figure 6D). Therefore, all four compounds appear to be specific inhibitors of nsp14 methyltransferase.

**Figure 6:**
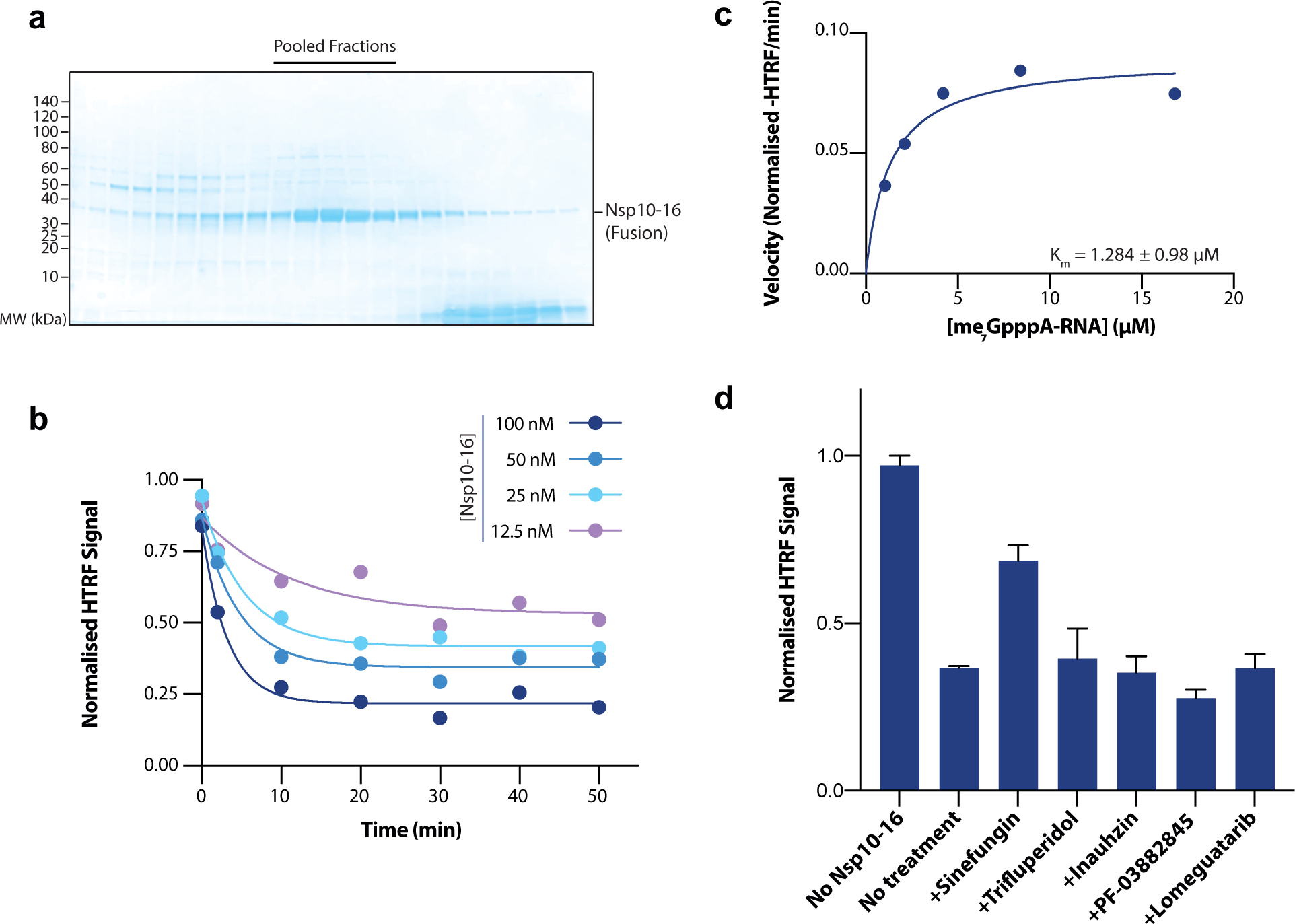
Nsp14 inhibitors do not inhibit nsp10-16. **a)** Gel filtration from final purification step of the nsp10-16 fusion protein. Coomassie shows fractions taken across the major peak of the gel filtration elution. Only fractions indicated were pooled (black bar, upper). Expected size of nsp10-16: 47.8 kDa (nsp10 - 13.3 kDa + nsp16 - kDa 34.5 kDa) **b)** Time course of the nsp10-16 methyltransferase reaction with 100 nM, 50 nM, 25 nM and 12.5 nM nsp10-16 enzyme. Reaction was conducted with 1.3 μM me_7_GpppA-RNA and 1 μM SAM. **c)** Determination of Michaelis constants for me_7_GpppA-RNA for nsp10-16. nsp10-16 concentration was fixed at 100 nM and substrate concentration varied. Errors given are 95% confidence ranges. **d)** Cross validation of nsp14 inhibitors with nsp10-16. Normalised HTRF values for nsp10-16 with inhibitors identified for nsp14 and sinefungin. All compounds tested at 50 μM. Reactions conducted with 1.3 μM me_7_GpppA-RNA and 1 μM SAM.

### Nsp14 inhibitors show antiviral activity in cellular infection models

In order to check if our compounds identified *in vitro* had any inhibitory effects on viral replication in mammalian cells, we utilised a SARS-CoV-2 viral infection assay using VERO E6 cells that are a model cell line for viral infection assays. We assayed viral replicative capacity by infecting cells with a constant amount of SARS-CoV-2 in the presence of inhibitor. We then quantified viral replicative capacity in fixed cells 22 hours post-infection using a fluorescent anti-Nucleoprotein antibody. This method allows the co-determination of cell viability through DNA staining, and ensures that our compounds were not reducing viral load through cytotoxicity rather than inhibiting viral replication (Figure 7A).

**Figure 7:**
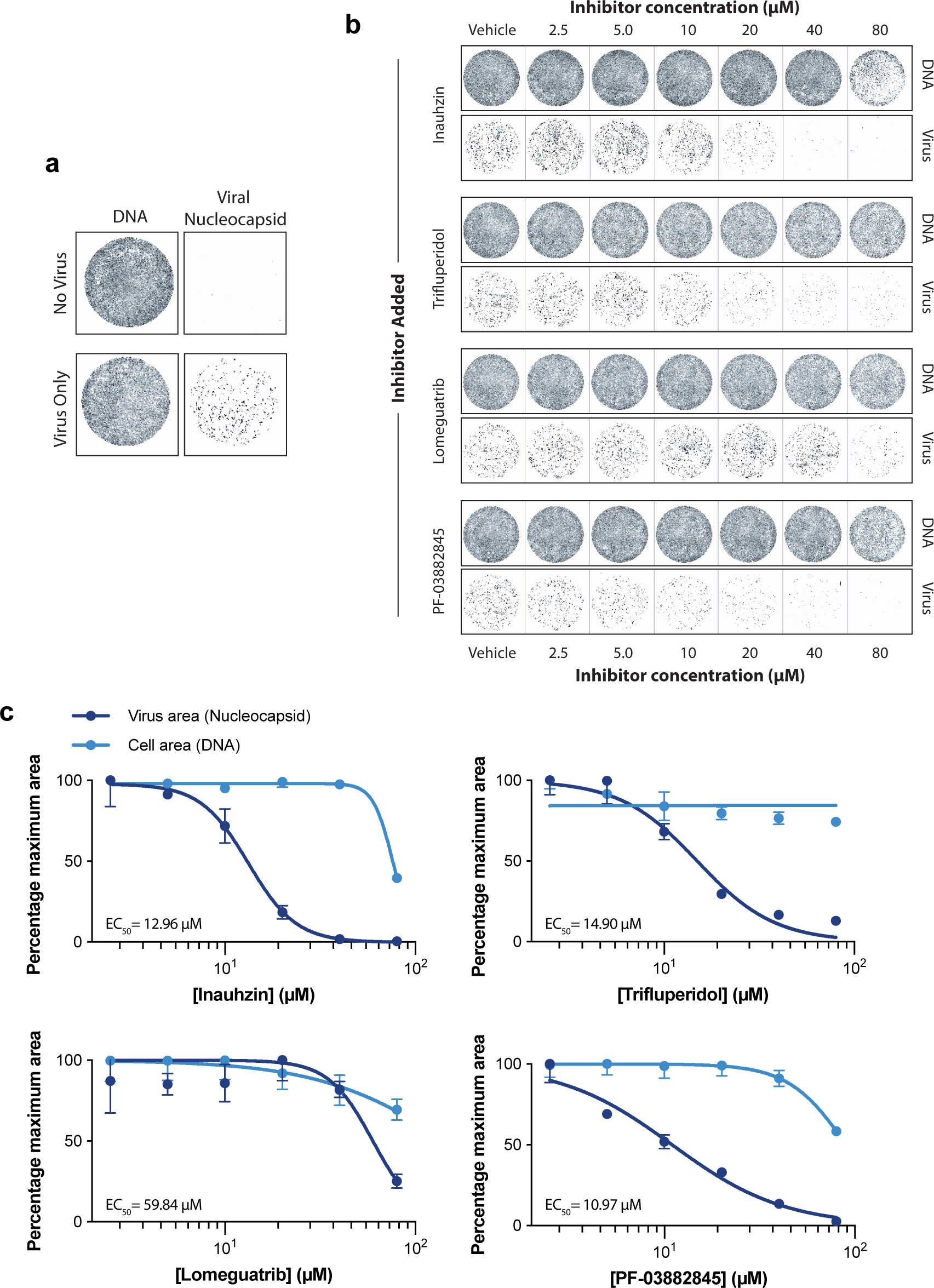
Nsp14 inhibitors are effective in cellular infection models. **a)** Representative wells showing VERO E6 cells stained for DNA using DRAQ7, and viral nucleoprotein using Alexa 488 conjugated anti-nucleocapsid antibody (see methods). Top panel: Cells with no virus added. Lower panel: Cells with virus added to a MOI of 0.5 PFU/cell. **b)** After seeding cells, media was washed and replaced with fresh media containing compounds at indicated concentrations, followed by infection with SARS-CoV-2. Wells shown are representative of three biological repeats. DNA is visualised through staining by DRAQ7, and virus is visualised through staining for viral nucleoprotein (see methods). **c)** Quantification of viral and cell area through fluorescence microscopy of DNA staining by DRAQ7 and viral nucleoprotein (see methods). DNA and viral area measurements were initially normalised to the vehicle control. Values were then plotted as a percentage of the maximum normalised measurement. IC_50_ values are given for each compound in the respective panel. Points represent mean values; error bars give standard deviation over three biological repeats. Error bars are not given if the error is smaller than the size of the point.

All four compounds showed antiviral activity at or below 80 μM, with limited cytotoxic effects (Figure 7B). The most effective compounds were PF-03882845 (EC_50_ = 10.97 μM), Inauhzin (EC_50_ = 12.96 μM) and Trifluperidol (EC_50_ = 14.9 μM). Lomeguatrib was less effective at inhibiting viral replication, with an EC_50_ = 59.84 μM. Concentrations at which Lomeguatrib was effective also came with slight cytotoxicity, whereas all three of PF-03882845, Inauhzin, and Trifluperidol show little to no cytotoxicity at their EC_50_ concentrations (Figure 7C). This establishes that the compounds we identified *in vitro* have antiviral activity in mammalian cells with limited cytotoxic effects.

At the moment, remdesivir is the only antiviral compound that can be taken both prophylactically and therapeutically for the treatment of COVID-19. Other treatments for COVID-19 are solely therapeutics that either modulate the immune response (such as dexamethasone) or are antibody-based therapeutics (such as bamlanivimab). Combination therapy, the use of two or more drugs with different modes of action, is a tested therapeutic strategy for the treatment of some diseases. In addition to achieving better physiological outcomes, combination therapies may also be an effective strategy for limiting antiviral drug resistance [27]. Therefore, we wished to see if the identified compounds from our screen demonstrated synergistic effects with remdesivir, which might warrant exploration for combination therapies and prophylaxis.

Although remdesivir is effective at inhibiting SARS-CoV-2 replication in cellular models, it is known that remdesivir has limited capacity to reduce viral titre in VERO E6 cells at 1 μM and below (Figure 8A) (25). Therefore, we conducted similar viral infection assays as described previously, but in additional presence of remdesivir at a concentration of 0.5 μM (Figure 8B).

**Figure 8:**
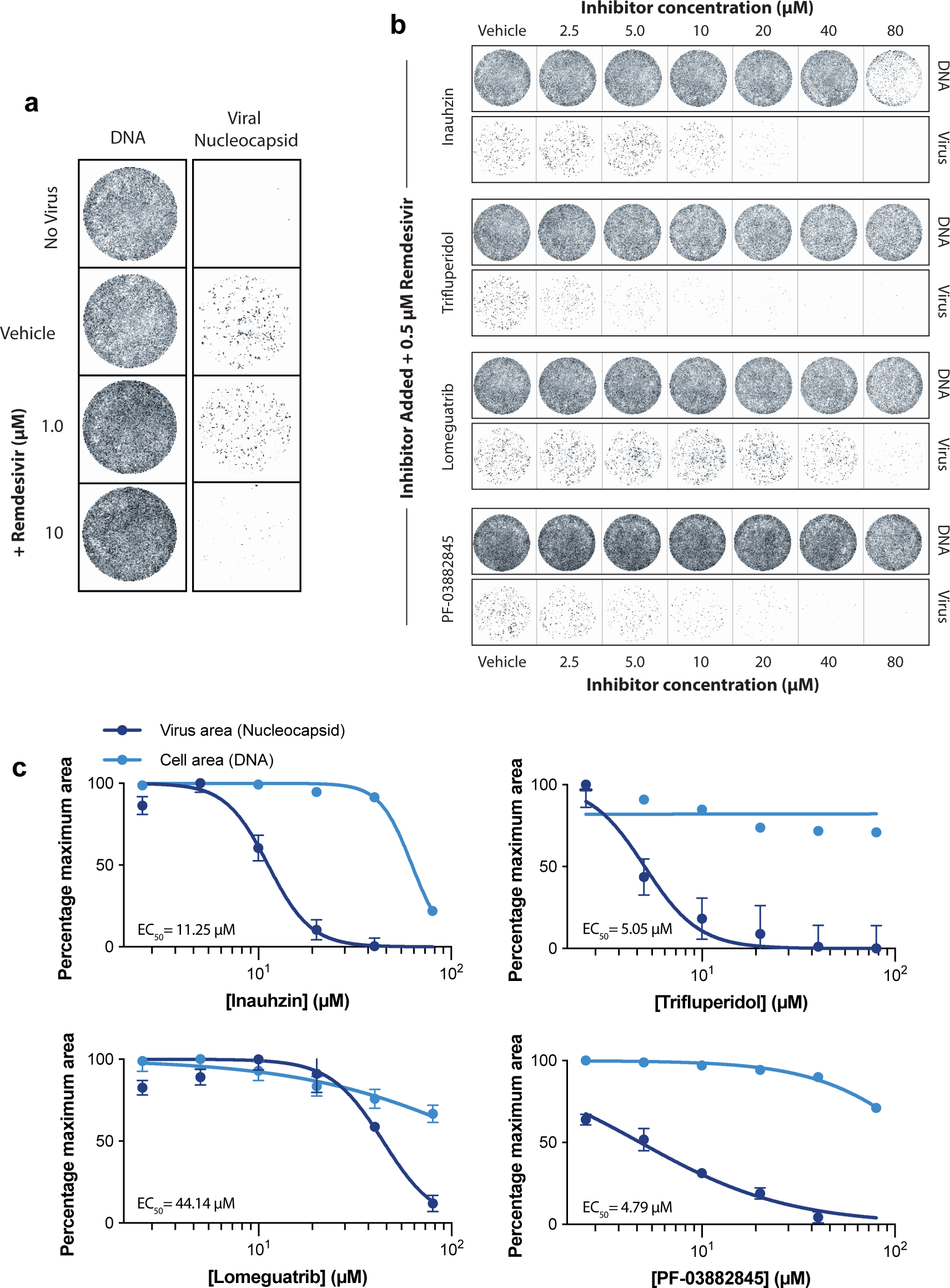
Nsp14 inhibitors are synergistic with remdesivir. **a)** Representative wells showing cells after no infection, or infection with remdesivir or vehicle control. DNA is visualised through staining by DRAQ7, and virus is visualised through staining for viral nucleoprotein (see methods). It is known that < 1 μM remdesivir is ineffective in VERO E6 Cells (25). **b)** After seeding cells, media was washed and replaced with fresh media containing compounds at indicated concentrations, followed by infection with SARS-CoV-2. In addition, 0.5 μM remdesivir was added to all wells. Wells shown are representative of three biological repeats. DNA is visualised through staining by DRAQ7, and virus is visualised through staining for viral nucleoprotein (see methods). **c)** Quantification of viral and cell area through fluorescence microscopy of DNA staining by DRAQ7 and viral nucleoprotein (see methods). DNA and viral area measurements were initially normalised to the vehicle control. Values were then plotted as a percentage of the maximum normalised measurement. IC_50_ values are given for each compound in the presence of remdesivir. Points represent mean values; error bars give standard deviation over three biological repeats. Error bars are not given if the error is smaller than the size of the point.

Inauhzin demonstrated similar IC_50_ values in the presence of remdesivir (Inauhzin EC_50_+Rem = 11.25 μM) (Figure 8C). However, PF-03882845, Trifluperidol, and Lomeguatrib showed markedly reduced IC_50_ values in the presence of remdesivir, with values falling close to a factor of 3 for the two former compounds (PF-03882845 EC_50_+Rem. = 4.79 μM; Trifluperidol EC_50_+Rem.= 5.05 μM Lomeguatrib EC_50_+Rem. = 44.14 μM), indicating a potential synergy between our identified nsp14 inhibitors and remdesivir (Figure 8C).

## Discussion

The SARS-CoV-2 RNA methyltransferases have been somewhat overlooked as a therapeutic target, currently with no characterised inhibitors *in vitro* or *in vivo*. Here we describe the purification and characterisation of both RNA cap methyltransferases encoded by the SARS-CoV-2 genome. In addition, we describe the successful discovery of novel inhibitors of nsp14 methyltransferase activity *in vitro*, which are not only effective in cell-based assays at low micromolar concentration, but demonstrate synergy with the only approved SARS-CoV-2 therapeutic, remdesivir. We show for the first time that SARS-Cov-2 nsp14 *in vitro* inhibitors are effective at cessation of viral replication, strongly suggesting that the methyltransferase activity of nsp14 is essential for coronavirus replication.

We initially demonstrated that a commercially available HTRF assay was able to detect the methyltransferase activity of both nsp14 and an nsp10-16 fusion protein. This level of quantitation allowed us to accurately determine the K_m_ values of nsp14 against: GTP, free GpppA, and GpppA-capped RNA; as well as the K_m_ for me_7_GpppA-RNA for nsp10-16. This revealed that whilst nsp14 preserved the ability to methylate guanine without need for the complete cap structure or attached RNA, that the K_m_ values for these substrates are significantly (∼100x) higher than the K_m_ of me_7_GpppA-RNA for nsp10-16. Thus, it appears that nsp14 is able to methylate a broad variety of substrates at the expense of a higher K_m_, whereas nsp10-16 has a lower K_m_ but is known to only methylate capped RNAs [28].

We identified and validated four antiviral compounds that were potential inhibitors of nsp14 methyltransferase activity. Of these compounds, Lomeguatrib and PF-03882845 have previously been evaluated in phase II trials for the treatment of melanoma and diabetic nephropathy, respectively [29, 30]. Lomeguatrib is an inhibitor of the O-6’-methylguanine-DNA methyltransferase [31], MGMT (IC_50_ ∼ 5 nM), whereas PF-03882845 is a mineralocorticoid receptor agonist (IC_50_ ∼ 10 nM) [32]. Although neither compounds were carried forward for phase III trials, they may be safe for human use. Trifluperidol is a licenced therapeutic currently in use for the treatment of psychoses including schizophrenia but has relatively severe side effects that limit its potential use as a prophylactic treatment for COVID-19, but may warrant its exploration as a post-infection antiviral. Finally, Inauhzin has previously been characterised as an inhibitor of SIRT1 (IC_50_ ∼ 1 μM) [33], but has not yet been taken forward into human trials. Although all are able to inhibit nsp14, these compounds have no obvious common chemical similarities (Figure S3), raising the potential of multiple inhibitory binding modes that may be exploited to generate future, more potent, nsp14 inhibitors.

The potency of inhibitors in mammalian cells was similar to the *in vitro* inhibition of Nsp14 for Lomeguatrib and Trifluperidol. The efficacy of PF-03882845 was significantly reduced in cells compared to the biochemical assays, which might suggest issues with cell permeability or that the drug is actively metabolised into a non-inhibitory form within cells. Inauhzin presented an interesting result as it appeared to be more potent in cells than in the biochemical assays. Perhaps Inauhzin, as a sirtuin inhibitor, for example, has other cellular effects aside from the inhibition of Nsp14 that may contribute to a reduction in viral infectivity. Although we cannot rule out that inhibition of other cellular pathways might be contributing to the observed reduction of viral load in cells, aside from the inhibition of Nsp14, we note that the IC_50_ values for the known functions of Lomeguatrib, Trifluperidol, and PF-03882845 are far below the IC_50_ values for viral load reduction. Therefore, for these three compounds, it is likely that they exert their effects on reduction of viral load through the inhibition of Nsp14.

As remdesivir currently stands as the only licenced antiviral for SARS-CoV-2 infection, we also tested to see if the nsp14 inhibitors we identified had any synergistic effect with remdesivir in reducing viral load. This combination treatment approach is commonly used to treat viral disease, most prominently in the treatment of HIV [34]. Among the four compounds, three showed synergy with remdesivir (Lomeguatrib, IC_50_ reduction∼27%; PF-03884528, IC_50_ reduction 56%; Trifluperidol IC_50_ reduction 66%). Only Inauhzin showed no synergy with remdesivir treatment, perhaps because it exerts an effect via other cellular targets. The IC_50_ of Trifluperidol and PF-03884528 was reduced in combination with remdesivir to a point where they may be clinically relevant. Given that both are apparently tolerated in humans, using them in combination with remdesivir could be a promising path for therapeutic exploration.

## Experimental Procedures

### Protein Expression and Purification

#### Nsp14

Rosetta™ (DE3) pLysS cells (Novagen) (F^-^ *ompT hsdS*_B_(r_B_^-^ m_B_^-^) *gal dcm* (DE3) pLysSRARE (Cam^R^) were transformed with plasmids expressing His_14_-SUMO-Nsp14 or His_14_-SUMO-Nsp14-D331A (DU70487 and DU70488; available from https://mrcppu-covid.bio/).

Transformant colonies were inoculated into a 100 ml LB / chloramphenicol (35 µg/ml) culture (containing kanamycin (50 µg/ml) and grown overnight at 37°C with shaking at 200 rpm.

Next morning, the culture was mixed with 400 ml of LB / chloramphenicol and kanamycin and further grown until OD_600_ reached 0.8. The protein expression was induced by 0.05 mM IPTG addition and the culture was shaken overnight at 18 °C.

Cells were harvested by centrifugation at 5000 rpm for 10 min in an JLA-9.1000 rotor (Beckman). The bacterial pellet was resuspended in 20 ml lysis buffer (50 mM Bis-Tris-HCl (pH 6.8), 0.5 M NaCl, 4 mM MgCl_2,_ 30 mM imidazole, 0.5 mM TCEP, Roche protease inhibitor tablets) with 500 µg / ml Lysozyme, then incubated at 4 °C for 0.5 h with rotation. Subsequently, the sample was sonicated twice for 90 s (15 s on, 30 s off) at 40% on a Branson Digital Sonifier. After centrifugation at 15,000 rpm at 4°C for 0.5 h in an JA-30.50 rotor (Beckman), the obtained soluble extract was mixed with 2 ml slurry Ni-NTA beads (30210, QIAGEN), incubated at 4 °C for 2 h with rotation.

Beads were recovered in a disposable gravity flow column and washed with 150 ml of lysis buffer then 20 ml lysis buffer containing 10 mM MgCl_2_ and 2 mM ATP (to remove bacterial chaperones) then 20 ml lysis buffer. Proteins were eluted with 4 ml lysis buffer containing 0.4 M imidazole. Subsequently, Ulp1 protease (10 mg/ml [35]) was added to cleave the HIS_14_-SUMO tag, incubated overnight at 4 °C. Untagged nsp14 was loaded onto a 120 ml Superdex 200 column in Gel filtration buffer (50 mM Bis-Tris-HCl (pH 6.8), 0.15 M NaCl, 4 mM MgCl_2_, 0.5 mM TCEP). nsp14-containing fractions were pooled, concentrated to 2.5 mg/ml by ultrafiltration using Amicon Ultra centrifugal unit (30 k MWCO; MERCK), aliquoted and snap frozen.

#### Nsp10-16

His-Sumo nsp10-16 fusion constructs were expressed in T7 express lysY/I^q^ *E. coli* cell (NEB). Cells were grown at 37°C to log phase to achieve OD 0.8. Cells were then induced by the addition of 0.5 mM IPTG and continued to incubate at 18°C overnight. Cells were harvested and lysed with in buffer A (50 mM HEPES-KOH, pH7.6, 10% glycerol, 1mM DTT, 0.02% NP-40, 300 mM NaCl and 30 mM imidazole), with addition of 100 ug/ml lyzsozyme and sonicate 24x5s. Lysate was centrifuged and supernatant was collected. The supernatant was incubated with Ni-NTA agarose beads (Thermo) for 1 hr at 4°C. Beads were wash with wash buffer A. The protein was eluted with 400 mM of imidazole. Fractions were pooled and dialyzed in buffer B (50 mM HEPES-KOH, pH7.6, 10% glycerol, 1mM DTT, 0.02% NP-40 and 150 mM NaCl) and 0.02 mg/ml His-Ulp1 to cleave-off the His-Sumo-tag. The dialysis was collected and incubate with Ni-NTA agarose beads once again to remove the proteases. The flow through was collected and load on Superdex S200 Increase 10/300 GL (GE healthcare) with buffer B. Peak fractions were collected and pooled.

#### Substrate generation for nsp14

nsp14 utilises GpppA capped RNA as a substrate. In order to synthesise this substrate RNA containing the first 30 nt of the SARS-CoV-2 genome (followed by 60 nt of junk RNA) was synthesised by firstly annealing Oligo_L3 (CAGTAATACGACTCACTATTa<u>G</u>taaggtttataccttcccaggtaacaacttgcttacccgaatcctatgaatttcctacgtcgtatctc) and Oligo_L3_Rev (gagatacgacgtaggaaattcataggattcgggtaagcaagttgttacctgggaaggtataaacctta<u>C</u>tAATAGTGAGT

CGTATTACTG). One A->G mutation was made in the 30nt region encoding the SARS-CoV-2 genome (underlined) to conform to improve transcription by T7 RNA polymerase. The DNA oligos were annealed in buffer containing 10 mM Tris-HCl, pH 7.5, 50 mM NaCl and 1 mM EDTA, heating to 95 °C, before cooling at 1 °C/minute until room temperature. RNA was co-transcriptionally capped with GpppA cap analogue (New England Biolabs) by using the HiScribe T7 High Yield RNA Synthesis (NEB) using all nucleotide triphosphates at 10 mM except for GTP, which was used at 0.75 mM. GpppA cap analogue was used at 9.25 mM. The reaction was run at 37 °C for 16 hrs, and purified using Monarch RNA cleanup kits (NEB).

#### Substrate generation for nsp16

Oligo_L3_ and oligo_L3_Rev were again annealed in buffer containing 10 mM Tris-HCl, pH 7.5, 50 mM NaCl and 1 mM EDTA. The template was synthesized using HiScribe t7 Quick High Yield RNA Synthesis (NEB) at 37 °C for 16 hrs, using all nucleotide triphosphates at 10 mM without any nucleotide substitutions. The synthesized RNA was purified using Monarch RNA cleanup kits (NEB). Purified RNA was then heated at 65 °C for 5 min to denature the secondary structure. Purified and denatured RNA was capped using Vaccinia Capping Kit (NEB) and finally purified using Monarch RNA cleanup kit.

### Assays

#### TLC assay for nsp14 enzymatic activity

The N-7 cap guanosine methylation assay was performed according to Varshney *et al.* [36]. Briefly, 20ng N14 was incubated with 200nM SAM and a 32P-G-capped substrate at 30C for 5-30 mins. ^32^P-m7G-capped and G-capped substrates were cleaved with P1 nuclease and caps resolved by thin layer chromatography on PEI cellulose/0.4M Ammonium sulphate.

Caps were quantitated by phosphor-imaging.

#### Methyltransferase assay

The methyltransferase activity of nsp14 was assayed by the detection of released SAH from the methyltransferase reaction. Released SAH was detected through the use of the commercially available EPIgeneous™ methyltransferase kit (CisBio Bioassays). Individual kit reagents were reconstituted according to the manufacturer’s instruction. For screening, the methyltransferase reaction was conducted at room temperature in an 8 μl reaction volume with 10 nM nsp14, 1 μM Ultrapure SAM (CisBio), 0.14 mM GpppA RNA cap analogue (New England Biolabs) in reaction buffer consisting of HEPES-KOH pH 7.6, 150 mM NaCl, and 0.5 mM DTT. The reaction was started with the addition of nsp14 and was allowed to proceed for 20 minutes before quenching by the addition of 2 μl 5M NaCl to a final concentration of 1M.

Following quenching, 2 μl Detection Buffer 1 (CisBio) was immediately added to the reaction mixture. After 10 minutes, 4 μl of 16X SAH-d2 conjugate solution (CisBio) was added. 16X SAH-d2 was prepared by adding one part SAH-d2 to 15 parts Detection Buffer 2 (CisBio).

After 5 minutes, 4 μl of 1X α-SAH Tb Cryptate antibody solution was added to the reaction mixture. 1X α-SAH Tb Cryptate antibody solution was prepared by adding one part α-SAH Tb Cryptate antibody (CisBio) to 49 parts Detection Buffer 2 (CisBio).

Homogenous Time Resolved Fluorescence (HTRF) measurements were taken after 1 hour following α-SAH Tb Cryptate antibody addition on a Tecan Infinite M1000 Pro plate reader. Readings were taken with a lag time of 60 μs after excitation at λ=337nm. Readings were taken emission wavelengths of λ=665nm and λ=620nm. The experimental HTRF ratio (HTRF_exp_) was then calculated as ratio of emission intensities: λ=665/λ=620. To reach the normalised HTRF ratio, HTRF ratio measurements were also taken of wells without enzyme (E_0_) and without SAH-d2 (d2_0_), representing the maximum and minimum achievable HTRF values, respectively. The normalised HTRF ratio was then calculated as a linear transformation of the experimental HTRF ratio, the E_0_ ratio, and the d2_0_ ratio:

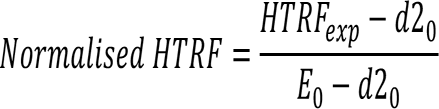

The buffer used for the nsp16 assay was 40 mM HEPES-KOH, pH7.6, 1 mM DTT, 5 mM MgCl_2_, 10% glycerol, 0.02% Tween-20, 1.3 mM substrate and 100 nM enzyme, unless stated otherwise. The inhibitor was pre-incubated with enzyme for 10 min at RT. EPIgenous Methyltransferase kit (Cisbio Bioassays) was used for the detection of methyltransferase activity as with nsp14.

#### High-throughput screening assay for nsp14 and hit validation

High-throughput screening was performed using a custom compound collection assembled from commercial sources (Sigma, Selleck, Enzo, Tocris, Calbiochem, and Symansis). 2.5 nl of a 10mM stock of the compounds dissolved in DMSO were arrayed and pre-dispensed into the assay plates using an Echo 550 (Labcyte), before being sealed and stored at -80C until screening day. When selecting ‘hit’ compounds, we removed aberrant high Z-scores due to screening errors. These screening errors arose due to systematic dispensation errors, and occurred within the same repeated wells, or cluster of wells within the screen. These wells were either discarded completely from all plates, or hits within these wells were discarded after the initial Z-score cut-off. This resulted in approximately 1% data loss. The initial Z’ factor to assess screen quality was conducted including these outlier wells, and therefore provides a conservative estimate of screen quality.

#### Viral inhibition assay

1.5x10^3^ Vero E6 cells (NIBC, UK) resuspended in DMEM containing 10% FBS were seeded into each well of 96-well imaging plates (Greiner 655090) and cultured overnight at 37C and 5% CO2. The next day, a 5x solution of compounds were generated by dispensing 10mM stocks of compounds into a v-bottom 96-well plate (Thermo 249946) and back filling with DMSO to equalise the DMSO concentration in all wells using an Echo 550 (Labcyte) before resuspending in DMEM containing 10% FBS. The assay plates with seeded VERO cells had the media replaced with 60µl of fresh growth media, then 20µl of the 5x compounds were stamped into the wells of the assay plates using a Biomek Fx automated liquid handler.

Finally, the cells were infected by adding 20µl of SARS-CoV2 with a final MOI of 0.5 PFU/cell. 22h post infection, cells were fixed, permeabilised, and stained for SARS-CoV2 N protein using Alexa488-labelled-CR3009 antibody produced custom (see section for Recombinant mAb production) and cellular DNA using DRAQ7 (ABCAM). Whole-well imaging at 5x was carried out using an Opera Phenix (Perkin Elmer) and fluorescent areas and intensity calculated using the Phenix-associated software Harmony (Perkin Elmer).

#### Recombinant mAb production

Heavy and light chain variable regions for CR3009, were synthesized (Genewiz) based on the GenBank sequences with regions of overlap to restriction digested human IgG1 vectors for assembly cloning (NEB) to produce plasmids: CR3009HC and CR3001KC. N-protein specific mAb CR3009 was produced by co-transfecting Expi293F cells (Life Technologies) in suspension growing at 37oC in 8% CO2 atmosphere in FreeStyle 293T medium (Life Technologies) with the plasmids. The supernatants were harvested 6-8 days post-transfection. as per the original study describing this mAb (van den Brink et al., 2005). The CR3009 Mab were purified by affinity chromatography using a 5 ml Protein G column (Cytiva) attached to an AKTA Pure system. Upon loading, the column was washed with PBS and bound Mabs eluted with 0.1M glycine pH 2.2 and immediately neutralized with 1M Tris, pH 8.0. The mAb-containing fractions were pooled and subjected to Size Exclusion Chromatography using a Superdex 200 16/600 prep grade column. The purified mAb CR3009 was labelled with Alexa Fluor 488-NHS (Cat#1812 Jena Biosciences) according to the instructions from the manufacturer.

## Author Contributions

**Souradeep Basu**: Conceptualization, Methodology, Validation, Formal analysis, Investigation, Resources, Writing - Original Draft, Writing - Review & Editing, Visualisation. **Tiffany Mak:**Conceptualization, Methodology, Validation, Formal analysis, Investigation, Resources, Writing - Original Draft, Writing - Review & Editing. **Rachel Ulferts**: Methodology, investigation. **Mary Wu**: Methodology, resources, investigation. **Tom Deegan:** Methodology, resources, investigation, Writing - Review & Editing. **Ryo Fujisawa**: Methodology, resources, investigation, Writing - Review & Editing. **Kang Wei Tan**: Methodology, Investigation, Formal analysis, Validation, Resources, Writing - Review & Editing. **Chew Theng Lim**: Methodology, Investigation, Formal analysis, Validation, Resources, Writing - Review & Editing. **Clovis Basier**: Investigation. **Berta Canal**: Resources, Writing - Review & Editing. **Joseph F. Curran:** Resources**. Lucy Drury:** Investigation. **Allison W. McClure:** Resources, Writing - Review & Editing. **Emma L. Roberts:** Resources. **Florian Weissmann:** Resources. **Theresa U. Zeisner:** Investigation, Writing - Review & Editing. **Rupert Beale:** Supervision, project administration. **Victoria H. Cowling:** Supervision, project administration, investigation, resources, visualisation, formal analysis, writing - review & editing. **Michael Howell:** Supervision, project administration, writing - review & editing. **Karim Labib:** Supervision, project administration, writing - review & editing. **John F.X. Diffley:** Conceptualization, Supervision, Methodology, Project administration, Funding acquisition, Writing - Review & Editing.

## Acknowledgements

We are grateful to Anne Early, Tobias Fuller, and Jingkun Zeng for their assistance. This work was supported by the Francis Crick Institute, which receives its core funding from Cancer Research UK (FC001066), the UK Medical Research Council (FC001066), and the Wellcome Trust (FC001066). S.B., T.M., C.B., J.F.C, and E.L.R. were supported by the Lord Leonard and Lady Estelle Wolfson Foundation. T.U.Z. received funding from the Boehringer Ingelheim Fonds. B.C. and F.W. have received funding from the European Union’s Horizon 2020 research and innovation programme under the Marie Skłodowska-Curie grant agreement Nos 895786 and 844211. This work was also funded by the Medical Research Council (core grant MC_UU_12016/13 to K.L.), Cancer Research UK (Programme Grant C578/A24558 to K.L.) and the Wellcome Trust (reference 204678/Z/16/Z) for a Sir Henry Wellcome Postdoctoral Fellowship to T.D., and a Wellcome Trust Senior Investigator Award (106252/Z/14/Z) to J.F.X.D.

## Data Availability Statement

All data associated with this paper will be deposited in FigShare (https://figshare.com/).

## Supplemental Material

**Supplemental Figure 1.**
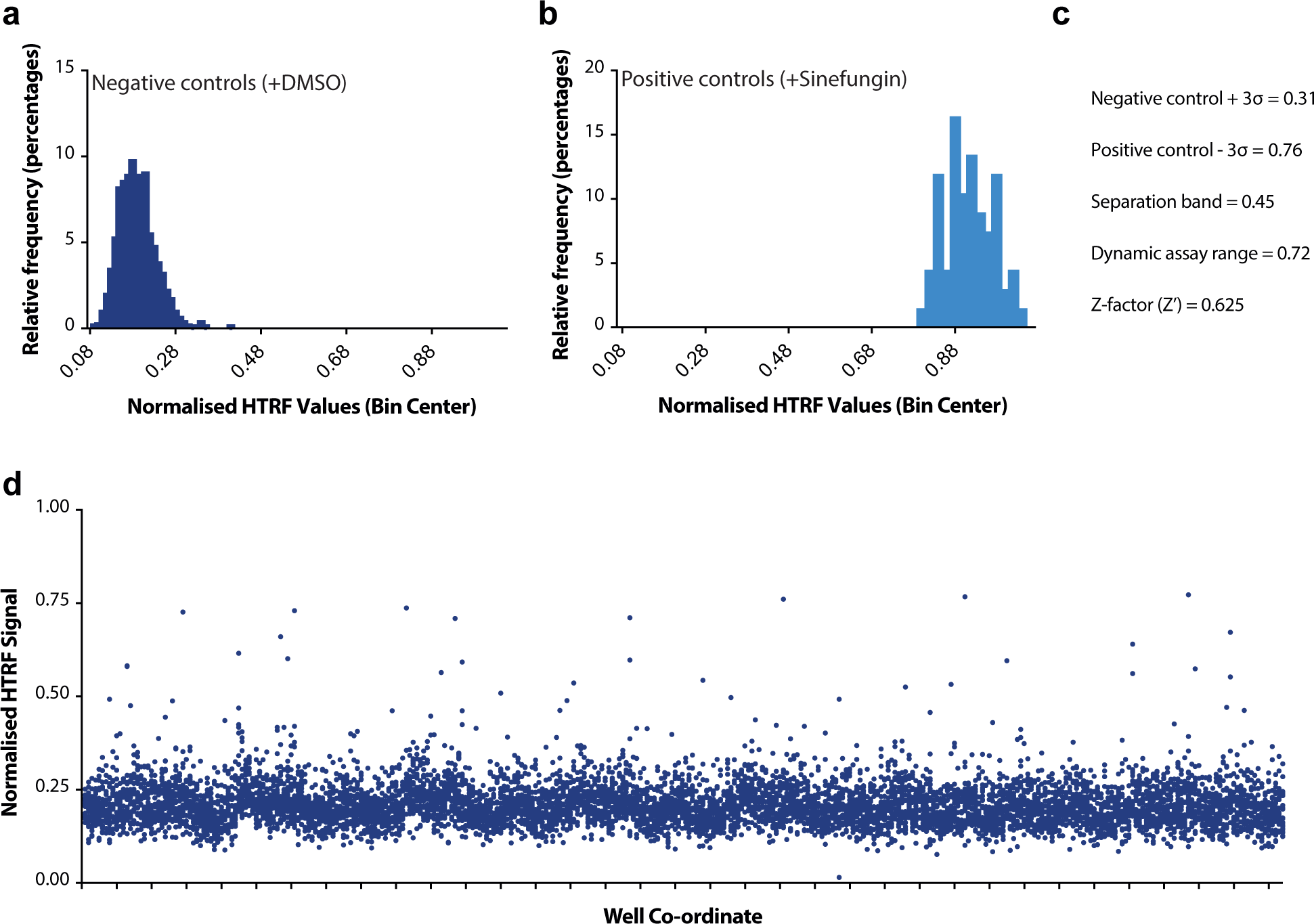
(S1)– Screening statistics. **a)** Normalised HTRF values for negative controls (+DMSO) within screen plates. N = 822. **b)** Normalised HTRF values for positive controls (+ Sinefungin) within screen plates. N = 67. **c)** Screening statistics as determined from the data in panels **(a)** and **(b)**. **d)** Raw normalised HTRF values over all compounds. Compounds are ordered by screening order, from first value read onwards.

**Supplemental Figure 2.**
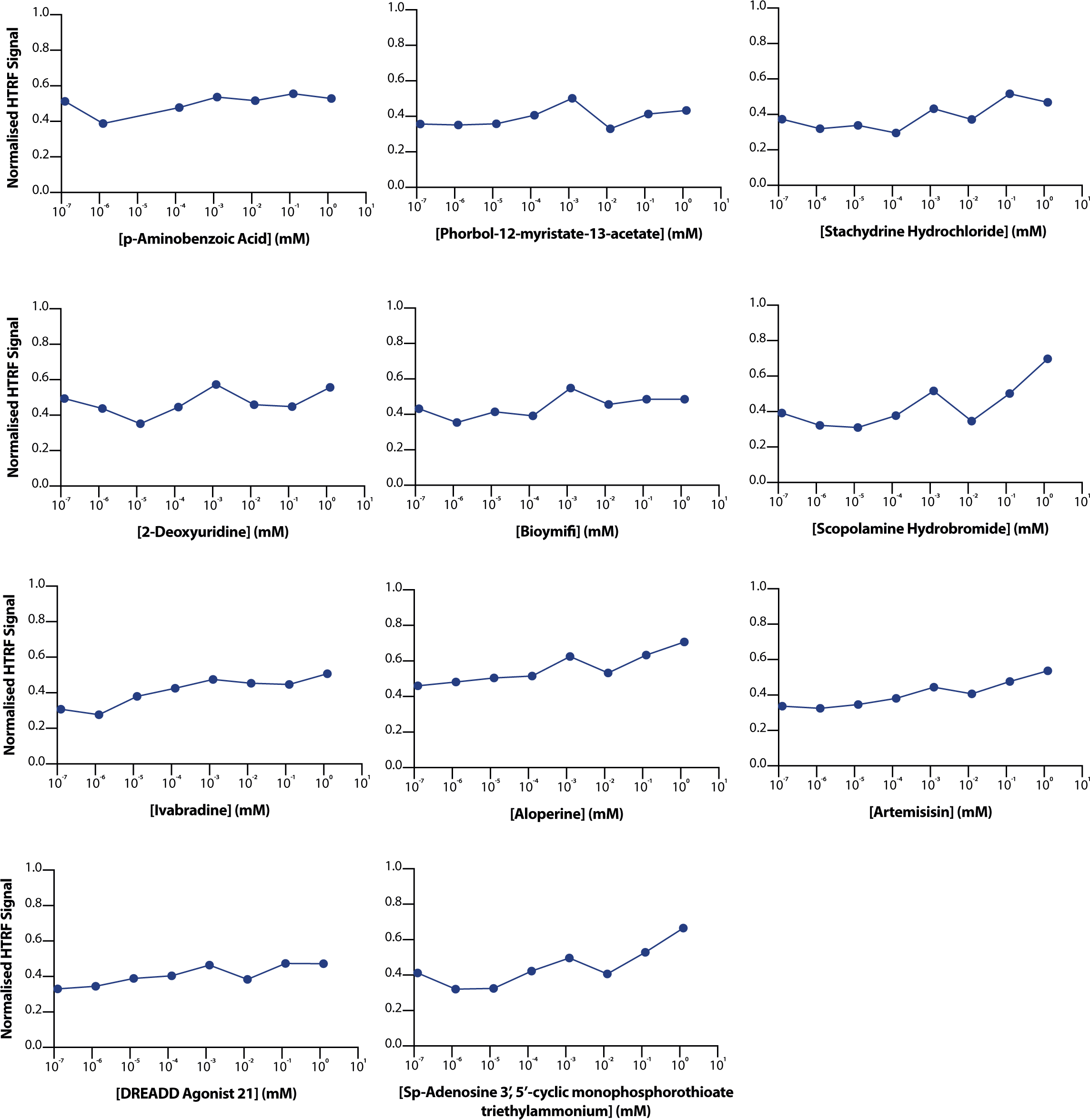
(S2)– Validation of other hit compounds. Normalised HTRF values for other screen hit compounds in validation. All reactions were run with 10 nM nsp14, 1 μM SAM, and 0.11 mM GpppA cap analogue for 20 minutes at room temperature before quenching with NaCl (see methods)

**Supplemental Figure 3.**
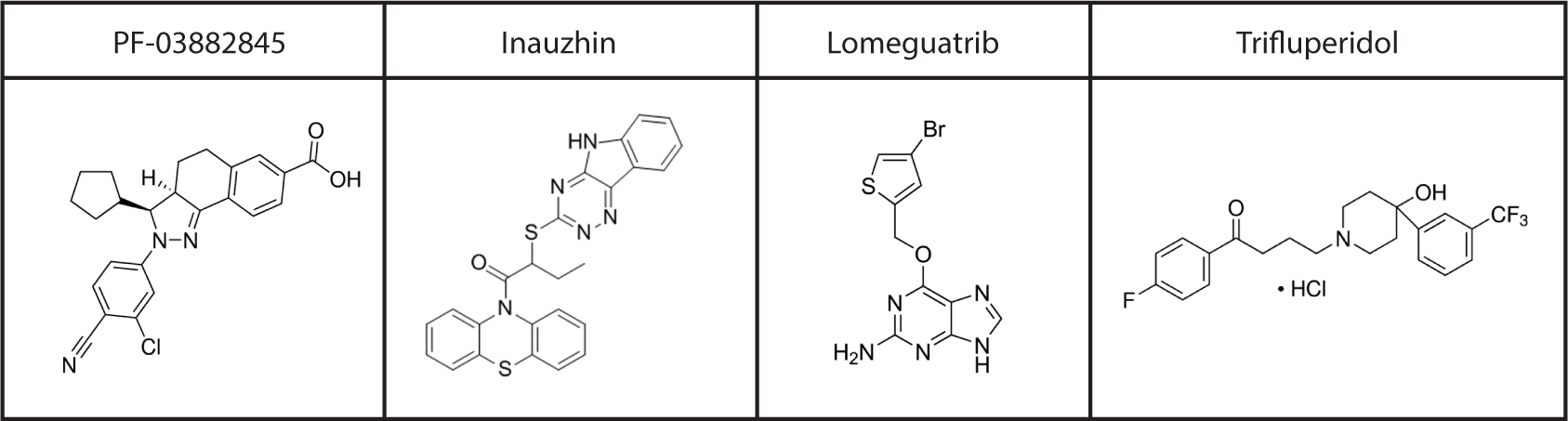
(S3)– Chemical structures of nsp14 inhibitors. Chemical structures of nsp14 inhibitors validated in HTRF assays and in cellular infection models.

Supplemental Table 1 (Table S1)– Hit compounds from screening List of 63 compounds that were considered “hits” from primary screening, and their associated Z-scores.

## References

1. Cucinotta, D. and M. Vanelli, WHO Declares COVID-19 a Pandemic. Acta Biomed, 2020. 91(1): p. 157–160.

2. Cui, J., F. Li, and Z.L. Shi, Origin and evolution of pathogenic coronaviruses. Nat Rev Microbiol, 2019. 17(3): p. 181–192.

3. Ortiz-Prado, E., et al., Clinical, molecular, and epidemiological characterization of the SARS-CoV-2 virus and the Coronavirus Disease 2019 (COVID-19), a comprehensive literature review. Diagn Microbiol Infect Dis, 2020. 98(1): p. 115094.

4. V’Kovski, P., et al., Coronavirus biology and replication: implications for SARS-CoV-2. Nat Rev Microbiol, 2021. 19(3): p. 155–170.

5. Singer, J.G., R.; Cotten, M.; Robertson, D., CoV-GLUE: A Web Application for Tracking SARS-CoV-2 Genomic Variation. Preprints, 2020.

6. Zhou, D., et al., Evidence of escape of SARS-CoV-2 variant B.1.351 from natural and vaccine-induced sera. Cell, 2021.

7. Finkel, Y., et al., The coding capacity of SARS-CoV-2. Nature, 2021. 589(7840): p. 125–130.

8. Ramanathan, A., G.B. Robb, and S.H. Chan, mRNA capping: biological functions and applications. Nucleic Acids Res, 2016. 44(16): p. 7511–26.

9. Furuichi, Y. and A.J. Shatkin, 5’-termini of reovirus mRNA: ability of viral cores to form caps post-transcriptionally. Virology, 1977. 77(2): p. 566–78.

10. Shuman, S., Structure, mechanism, and evolution of the mRNA capping apparatus. Prog Nucleic Acid Res Mol Biol, 2001. 66: p. 1–40.

11. Langberg, S.R. and B. Moss, Post-transcriptional modifications of mRNA. Purification and characterization of cap I and cap II RNA (nucleoside-2’-)- methyltransferases from HeLa cells. J Biol Chem, 1981. 256(19): p. 10054–60.

12. Case, J.B., et al., Mutagenesis of S-Adenosyl-l-Methionine-Binding Residues in Coronavirus nsp14 N7-Methyltransferase Demonstrates Differing Requirements for Genome Translation and Resistance to Innate Immunity. J Virol, 2016. 90(16): p. 7248–7256.

13. Cougot, N., S. Babajko, and B. Seraphin, Cytoplasmic foci are sites of mRNA decay in human cells. J Cell Biol, 2004. 165(1): p. 31–40.

14. Gu, M. and C.D. Lima, Processing the message: structural insights into capping and decapping mRNA. Curr Opin Struct Biol, 2005. 15(1): p. 99–106.

15. Kawai, T. and S. Akira, Innate immune recognition of viral infection. Nat Immunol, 2006. 7(2): p. 131–7.

16. Zust, R., et al., Ribose 2’-O-methylation provides a molecular signature for the distinction of self and non-self mRNA dependent on the RNA sensor Mda5. Nat Immunol, 2011. 12(2): p. 137–43.

17. Daffis, S., et al., 2’-O methylation of the viral mRNA cap evades host restriction by IFIT family members. Nature, 2010. 468(7322): p. 452–6.

18. Chen, Y., et al., Functional screen reveals SARS coronavirus nonstructural protein nsp14 as a novel cap N7 methyltransferase. Proc Natl Acad Sci U S A, 2009. 106(9): p. 3484–9.

19. Diamond, M.S., IFIT1: A dual sensor and effector molecule that detects non-2’-O methylated viral RNA and inhibits its translation. Cytokine Growth Factor Rev, 2014. 25(5): p. 543–50.

20. Menachery, V.D., K. Debbink, and R.S. Baric, Coronavirus non-structural protein 16: evasion, attenuation, and possible treatments. Virus Res, 2014. 194: p. 191–9.

21. Galloway, A. and V.H. Cowling, mRNA cap regulation in mammalian cell function and fate. Biochim Biophys Acta Gene Regul Mech, 2019. 1862(3): p. 270–279.

22. Bouvet, M., et al., RNA 3’-end mismatch excision by the severe acute respiratory syndrome coronavirus nonstructural protein nsp10/nsp14 exoribonuclease complex. Proc Natl Acad Sci U S A, 2012. 109(24): p. 9372–7.

23. Decroly, E., et al., Crystal structure and functional analysis of the SARS-coronavirus RNA cap 2’-O-methyltransferase nsp10/nsp16 complex. PLoS Pathog, 2011. 7(5): p. e1002059.

24. Stein, A., et al., Key steps in ERAD of luminal ER proteins reconstituted with purified components. Cell, 2014. 158(6): p. 1375–1388.

25. Jin, X., et al., Characterization of the guanine-N7 methyltransferase activity of coronavirus nsp14 on nucleotide GTP. Virus Res, 2013. 176(1-2): p. 45–52.

26. Ma, Y., et al., Structural basis and functional analysis of the SARS coronavirus nsp14-nsp10 complex. Proc Natl Acad Sci U S A, 2015. 112(30): p. 9436–41.

27. Al-Lazikani, B., U. Banerji, and P. Workman, Combinatorial drug therapy for cancer in the post-genomic era. Nat Biotechnol, 2012. 30(7): p. 679–92.

28. Chen, Y., et al., Biochemical and structural insights into the mechanisms of SARS coronavirus RNA ribose 2’-O-methylation by nsp16/nsp10 protein complex. PLoS Pathog, 2011. 7(10): p. e1002294.

29. Khan, O.A., et al., A phase II trial of lomeguatrib and temozolomide in metastatic colorectal cancer. Br J Cancer, 2008. 98(10): p. 1614–8.

30. A Study To Evaluate The Safety And Tolerability Of PF-03882845 In Patients With Type 2 Diabetic Nephropathy. 2012; Available from: https://ClinicalTrials.gov/show/NCT01488877.

31. Ranson, M., et al., Lomeguatrib, a potent inhibitor of O6-alkylguanine-DNA-alkyltransferase: phase I safety, pharmacodynamic, and pharmacokinetic trial and evaluation in combination with temozolomide in patients with advanced solid tumors. Clin Cancer Res, 2006. 12(5): p. 1577–84.

32. Orena, S., et al., PF-03882845, a non-steroidal mineralocorticoid receptor antagonist, prevents renal injury with reduced risk of hyperkalemia in an animal model of nephropathy. Front Pharmacol, 2013. 4: p. 115.

33. Zhang, Q., et al., A small molecule Inauhzin inhibits SIRT1 activity and suppresses tumour growth through activation of p53. EMBO Mol Med, 2012. 4(4): p. 298–312.

34. Cohen, M.S., et al., Prevention of HIV-1 infection with early antiretroviral therapy. N Engl J Med, 2011. 365(6): p. 493–505.

35. Deegan, T.D., et al., CMG helicase disassembly is controlled by replication fork DNA, replisome components and a ubiquitin threshold. Elife, 2020. 9.

36. Varshney, D., et al., Molecular basis of RNA guanine-7 methyltransferase (RNMT) activation by RAM. Nucleic Acids Res, 2016. 44(21): p. 10423–10436.

